# Cost-precision trade-off relation determines the optimal morphogen gradient for accurate biological pattern formation

**DOI:** 10.1101/2021.04.14.439772

**Authors:** Yonghyun Song, Changbong Hyeon

## Abstract

Spatial boundaries growing into macroscopic structures through animal development originate from the pre-patterning of tissues by signaling molecules, called morphogens. To establish accurate boundaries, the morphogen concentration which thresholds the expression of target gene at the boundary should be precise enough, exhibiting large gradient and small fluctuations. Producing more morphogens would better serve to shape more precise target boundaries; however, it incurs more thermodynamic cost. In the classical diffusion-degradation model of morphogen profile formation, the morphogens synthesized from a local source display an exponentially decaying concentration profile with a characteristic length λ. Our theory suggests that in order to attain a precise morphogen profile with the minimal cost, λ should be roughly half the distance to the target boundary position from the source, so that the boundary is formed at the position where the morphogen concentration is ∼10 % of the value at the source. Remarkably, we find that the well characterized morphogens that pattern the fruit fly embryo and wing imaginal disk form profiles with nearly optimal λ, which underscores the thermodynamic cost as a key physical constraint in the morphogen profile formation.

---

Complex spatial structures are shaped across the entire body from an ensemble of initially identical cells during animal development. The emergence of spatial structures is generally linked to the pre-patterning of tissues with biomolecules, called morphogens, that instruct cells to acquire distinct cell fates in a concentration dependent manner^1–3^. At the molecular level, the positional information encoded in the local morphogen concentration is translated by the cells into the expression of specific genes associated with cell-fate determination. In order to generate reproducible spatial organization, the concentration profile of morphogens must be stable against the noisy background signals inherent to cellular environment.

Along with quantitative measurements of morphogen gradients^4–10^, a number of theoretical studies have been devoted to understanding the precision and speed by which morphogen gradients are formed and interpreted^11–19^; however, generation of morphogen gradient, which breaks the spatial symmetry, incurs thermodynamic cost, and the relation of this cost with the precision and speed of morphogen gradient formation has rarely been addressed except for a few cases^11^. Creating and maintaining morphogen gradients require an influx of energy^20^, which is a limited resource for biological systems^21^, particularly at the stage of embryonic development^22,23^. In the present work, we study the trade-off between the cost and precision of the morphogen profile formation in the framework of the reaction-diffusion model of localized synthesis, diffusion, and depletion (SDD model)^4–6,24,25^.

The precision associated with morphogen gradients are perhaps best exemplified by the patterning of the anterior-posterior axis of the *Drosophila* embryo through the Bicoid (Bcd) gradient. In the two-hour post-fertilization of fruit fly embryogenesis, a uniform field of ∼6000 nuclei characterizes the periphery of the shared cytoplasm^26^. Bcd, a transcription factor, is produced from the anterior-end of the embryo by maternally deposited *bcd* mRNAs. Its subsequent diffusion and degradation engender an exponentially decaying profile of *bcd* concentration^27,28^, which is translated into the anterior expression of the target gene, *hunchback* (*hb*)^29^. Nuclei at around the middle section of the embryo can infer their relative spatial positions by detecting the concentration of Bcd with an exquisite precision of ∼ a single nucleus width^4^.

Other quantitatively characterized morphogens include Wingless (Wg), Hedgehog (Hh) and Decapentaplegic (Dpp), which pattern the dorsal-ventral (DV) and anterior-posterior (AP) axis of the wing imaginal disk in the fly larvae. Wg, a member of the Wnt signaling pathway, spreads through the wing disk from a narrow band of cells at the DV boundary^30,31^. The Wg concentration profile leads to the differential activation of multiple target genes, and, in particular, induces the sharp expression boundary of *senseless* (*sens*) a few cells away from the DV boundary^32,33^. Similarly, Hh patterns the AP axis by spreading from the posterior to the anterior side of the wing disk, affecting the expression of multiple genes. Its downstream transcriptional regulation gives rise to a strip of 8 – 10 *dpp* expressing cells that form the AP boundary^34,35^. Dpp, which spreads out from the localized production at the AP boundary, further patterns the AP axis. Major changes in the expression level of Dpp target genes, such as *spalt-major* (*salm*), occur at ~ 50 % of the total length of the domain patterned by Dpp^36,37^. Overall, *Drosophila* wing structures with the spatial precision of a single cell width emerge from the coordinated actions of multiple patterning events^38^.

The positional information, encoded in the concentration profile *ρ*(*x*) (red line in Fig. 1), is decoded to yield the target gene expression profile, *g*(*ρ*(*x*)) (blue line in Fig. 1). The morphogen profile, *ρ*(*x*), and the corresponding gene expression level, *g*(*ρ*(*x*)), together specify the cell fate in a morphogen concentration-dependent manner. In what follows, we will motivate a quantitative expression for the positional error associated with the “boundary” positions, where the target gene expression profile displays a sharp change.

**FIG. 1.**
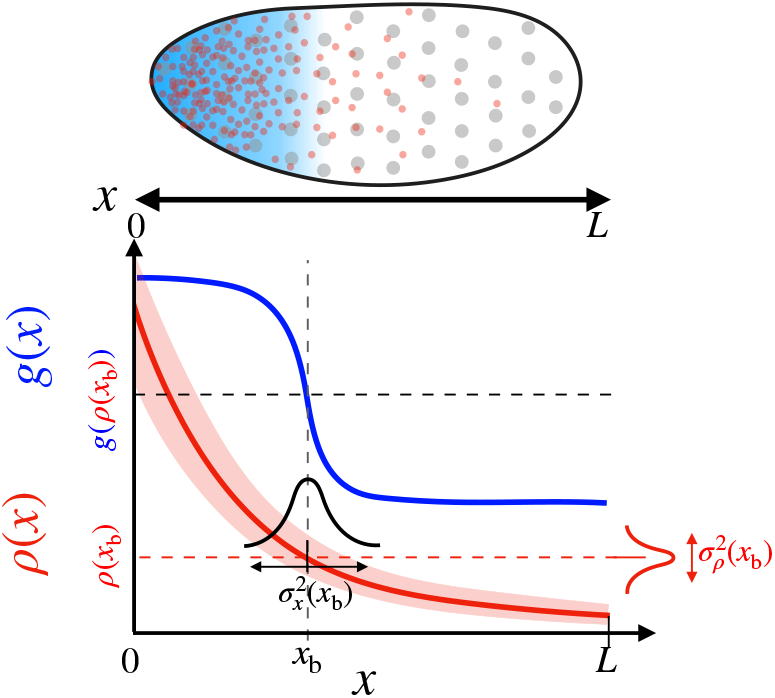
Positional information transfer by the morphogen gradient. (Top) The specification of the anterior region of the fruit fly embryo. The uniformly distributed nuclei (grey circles) are subjected to different levels of the morphogen (red dots) in the local environment, which leads to the anterior expression of the target gene (blue shade). (Bottom) The red and blue lines respectively depict the morphogen profile, ρ(x), and the target gene expression, g(x), which together specify cell fate. The squared positional error at the boundary x_b_, 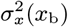, is defined as the product between the variance of the morphogen profile, 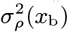, and the squared inverse slope of the morphogen profile, 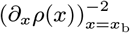.

We begin by defining a target boundary, *x* = *x*_b_, where the morphogen concentration is at its critical threshold value *ρ*_b_ ≡ *ρ*(*x*_b_). The nuclei exposed to morphogen concentrations higher than *ρ*_b_ would adopt an anterior cell fate. Prior to measuring the local morphogen concentration, each nucleus has no information regarding its own position (Fig. 1). After measuring 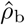, the nucleus can estimate the location of target boundary, such that 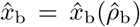. The error (variance) associated with the measured concentration at *x* = *x*_b_ with respect to its true value *ρ*_b_ is given by 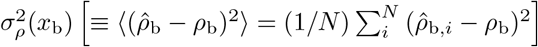.

Then, the Taylor expansion of the concentration at the target boundary, 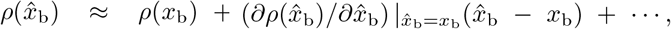, allows one to relate the error in measured concentration with the positional error in estimating the target boundary 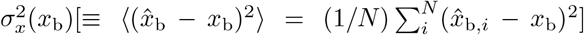 as follows:

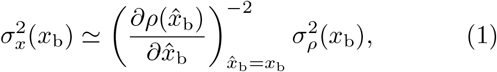

such that the morphogen profiles exhibiting large gradient and small fluctuations give rise to small positional errors.

Of fundamental importance is to address how naturally occurring morphogen profiles, tasked with the transfer of positional information, are formed under limited amount of resources. In an earlier work, by considering the total morphogen content at steady state as a proxy for the cost, Emberly evaluated the “cost-effectiveness” of the exponentially decaying Bcd profile with a characteristic decay length λ^11^. Remarkably, it was shown that for a given positional error, the Bcd profile is shaped with a cost-minimizing λ^11^. Here, we expand this argument with a in-depth treatment of the dynamics of morphogen profile formation and the thermodynamic cost involving the formation and maintenance of precise morphogen profiles.

## RESULTS

The cost of transferring the positional information by the steady state morphogen profile should include the cellular resources used to generate the profile prior to the measurement, as well as the resources to maintain the profile during the measurement. In this context, we consider two limiting scenarios. (i) Point measurement, in which the morphogen profile is measured instantaneously; (ii) Space-time-averaged measurement, in which the morphogen profile is measured over a finite space and for a time interval *T*. In the former, the associated cost is defined as the amount of morphogen produced while the profile approaches to the steady state. In the latter, we assume a space-time averaged measurement carried out for a long time duration. Then, the morphogen produced over the measurement can be approximated as the total cost required to create and maintain the morphogen gradient. For both of the limiting scenarios, the cost of generating morphogen profiles and the precision of the profiles are counterbalanced. We consider a *morphogen production-independent quantity* by taking the product of total cost and the squared relative error to quantify the trade-off between the cost and precision of the morphogen profile, and show that the trade-off product can be minimized when the morphogen gradient’s characteristic length, λ, is properly selected. We evaluate the cost-effectiveness of morphogen profiles patterning the fruit fly embryo and wing disk, which have been quantitatively characterized ^4–6,10,33,39–44^.

### Point measurement

We begin by defining the dynamics of morphogen profile formation in a system composed of a one-dimensional array of cells in the domain 0 ≤ *x* ≤ *L*, where *L* is the system size (see Figs. 1 and 2(a)). The term ‘cell’ refers to the spatial grid in which to define the local positional error and the cost. We denote the amount of morphogen in the cell of the volume (*v*_cell_) and length (*l*_cell_) at the interval between *x* and *x*+*l*_cell_ by *ρ*(*x, t*) [conc ≡#*/v*_cell_]. Unless otherwise specified, *ρ* refers to the ensemble averaged morphogen concentration. At the left boundary (see Fig.2a), the morphogen is injected at a constant flux of *j*_in_ [conc × *l*_cell_*/*time]. At all positions, the morphogens are depleted with rate *k*_d_ [time^−1^], while spreading across the cells with the diffusivity 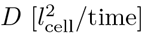 (Fig. 2a). For *l*_cell_*/L* ≪ 1 (i.e., at the continuum limit), the spatiotemporal dynamics of the morphogen is described using the reaction-diffusion equation

**FIG. 2.**
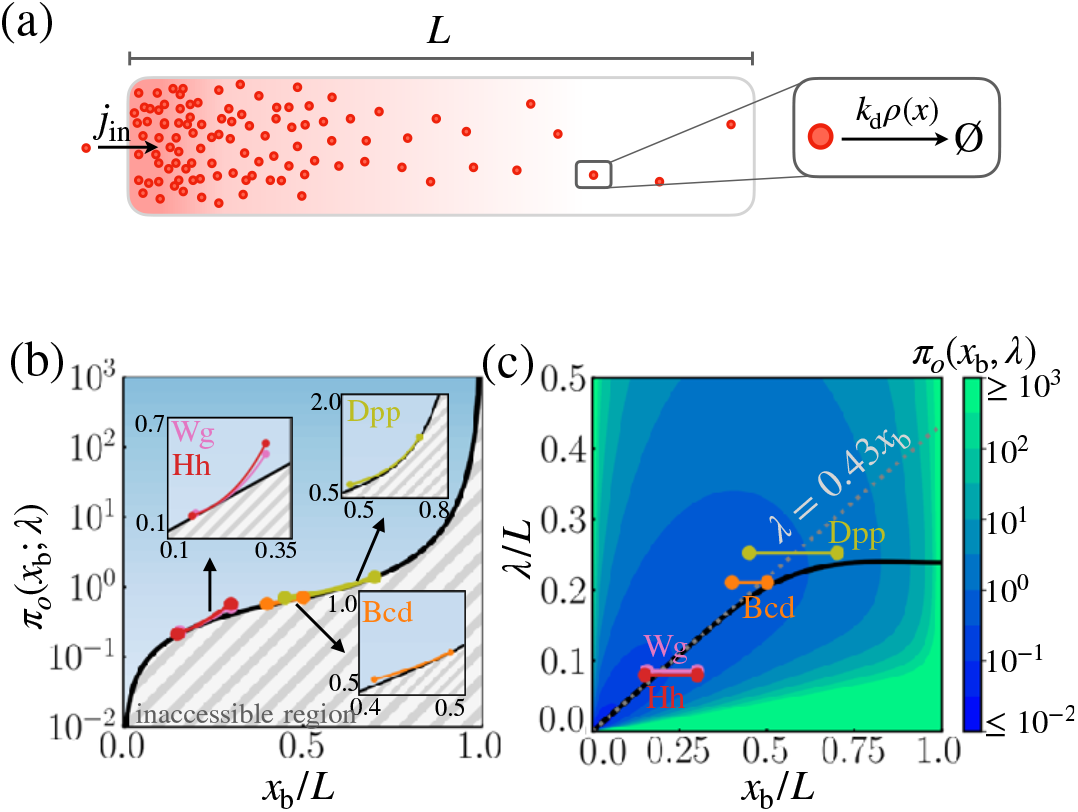
Cost-precision trade-off in the point measurement. **(a)** Schematic of the model. **(b)** The position-dependent lower bound of trade-off product *π*_*o*,min_(*x*_b_) obtained numerically. The grey hashed regions represent the inaccessible regions. The trade-off product of the morphogens profiles of Bcd, Wg, Hh, and Dpp are shown in the respective insets. **(c)** The black line denotes the optimal characteristic decay length (λ_min_) with respect to the position *x*_b_*/L*. The color scale indicates the trade-off product *π*_*o*_(*x*_b_) computed for each pair of λ*/L* and *x*_b_*/L* values. In **b** and **c**, the depicted trade-off product is normalized by *α*_*o*_*L/l*_cell_. The grey dotted line depicts the linear approximation of λ_min_ at large *L*. The parameters for the naturally occurring morphogen profiles are further described in the SI Appendix **Length scales of Bcd, Wg, Hh, and Dpp**.

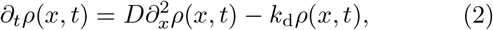

with boundary conditions −*D∂*_*x*_*ρ*(*x, t*) |_*x*=0_= *j*_in_ and *D∂*_*x*_*ρ*(*x, t*)|_*x*=*L*_ = 0. Then, the concentration profile at steady state is obtained as

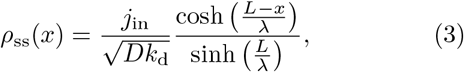

where 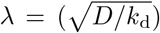 is the characteristic decay length of the concentration profile determined by the diffusivity of morphogen (*D*) and degradation rate (*k*_d_). In what follows, we will define (i) the *cost*, and (ii) the *precision* associated with this reaction-diffusion model.

(i) With a function *R*(*x, t*) ≡ (*ρ*_ss_(*x*) −*ρ*(*x, t*))*/*(*ρ*_ss_(*x*) −*ρ*(*x*, 0)), which captures the evolution of the morphogen profile from *ρ*(*x, t* = 0) = 0 to the steady state value *ρ*_ss_(*x*) (Eq. S2), the mean local accumulation time at *x* that characterizes the average time scale for establishing the steady state profile is calculated as^13^

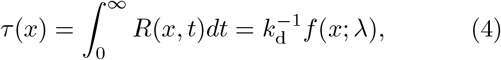

where 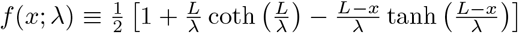 is a dimensionless quantity determined by *x* and λ. Then, *v*_cell_*j*_in_*/l*_cell_ × *τ* (*x*) quantifies the total number of morphogens produced over the time scale of *τ* (*x*). The thermodynamic cost of producing the morphogens, which is effectively the driving force of the pattern formation, is expected to be proportional to this number^45^, such that

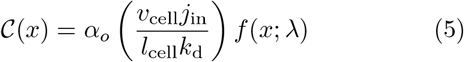

with a proportionality constant *α*_*o*_.

(ii) The local measure of precision at position *x* can be quantified by the squared relative error in the positional measurement (Eq. 1) as follows:

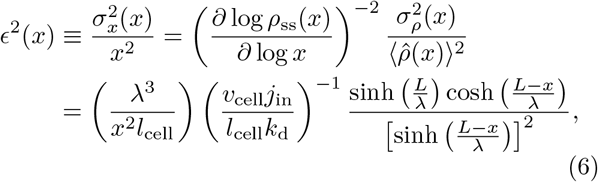

where 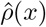 represents the morphogen concentration estimated at *x* from a measurement. Note that the multiple measurements of 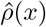 yield the mean concentration profile 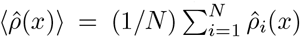, which is equivalent to *ρ*_ss_(x) at steady states given in Eq.S2. We use the fact that the probability distribution of the steady state concentration profile obeys the Poisson statistics^46^ (i.e.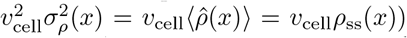, since all the reactions in the system are of zeroth or first order.

There is a trade-off between 𝒞(*x*) and *ϵ*^2^(*x*). If the *n*_*m*_ is the number of morphogen produced for a certain time duration, then 𝒞 (*x*) ∝ *n*_*m*_ and 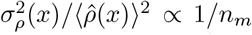, the latter of which simply arises from the central limit theorem, or can be rationalized based on the Berg-Purcell result 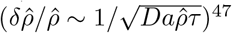 where *a* is the radius of volume in which a receptor detects the morphogen and *τ* is the detection time. Increasing the overall morphogen content reduces the morphogen profile’s positional error at the expense of a larger thermodynamic cost. In Eqs. 5 and 6, *n*_*m*_ corresponds to (*v*_cell_*j*_in_*/l*_cell_*k*_d_) (the morphogen produced for the time duration 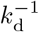), which cancels off when taking the product between (*x*) and *ϵ*^2^(*x*). Thus, the morphogen production-independent measure of trade-off between the cost of morphogen production and the squared relative error in position can be quantified by taking the product of the two quantities,

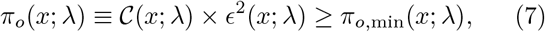

where the λ dependence of the trade-off product is made explicit for the discussion that follows.

We are mainly concerned with the precision at the boundary position, *x* = *x*_b_, where a sharp change in the downstream gene expression, *g*(*x*), is observed. The inequality Eq.7 constrains dynamical properties of the morphogen profile at *x*_b_, specifying either the minimal cost of morphogen production for a given positional error or the minimal error in the morphogen profile for a given thermodynamic cost (Fig. 2). In other words, when the trade-off product *π*_*o*_(*x*_b_; λ) is close to its lower bound *π*_*o*,min_(*x*_b_; λ), the system is cost-effective at generating precise morphogen profiles at *x*_b_. Generally, *π*_*o*,min_(*x*_b_; λ) increases monotonically with *x*_b_, which signifies that boundaries farther away from the origin require more morphogen production to achieve comparable positional error (Fig. 2b). For a fixed system size *L* and position *x*_b_, the value of the trade-off product is determined solely by the decay length, λ. By tuning λ, we can find the value of λ that minimizes the trade-off product at *x* = *x*_b_, i.e., *π*_*o*,min_(*x*_b_), such that

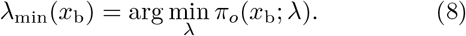

The optimal decay length, λ_min_(*x*_b_), also increases monotonically with the target boundary *x*_b_; for *x*_b_ *< L/*2, λ_min_ is well approximated by the linear relationship λ_min_(*x*_b_) ≈ 0.43*x*_b_ (See Eq. S28 for the derivation).

It is of great interest to compare *x*_b_ and λ of real morphogen profiles against the optimal λ_min_(*x*_b_). For Bcd, Wg, Hh, and Dpp morphogen profiles, we first estimated the possible range of *x*_b_ from the experimentally measured expression profiles of the respective target genes, *hb, sens, dpp* and *salm*^33,42,48^, *and then found that the characteristic lengths of the four morphogen profiles*, λ_Bcd_ = 100 *µ*m, λ_Wg_ = 6 *µ*m, λ_Hh_ = 8 *µ*m, and λ_Dpp_ = 20 *µ*m, are close to their respective optimal values, λ_min_(*x*_b_)^4,5^. Thus, our findings indicate that the concentration profiles of the four morphogens are effectively formed at the lower bound of the cost-precision trade-off, *π*_*o*,min_(*x*_b_) (Fig. 2b) (see the SI Appendix, **Length scales of Bcd, Wg, Hh, and Dpp** for more details on the relevant parameters of the morphogen profiles).

### Space-time-averaged measurement

The positional information available from the morphogen profile varies depending on the method by which the cell senses and interprets the local morphogen concentration. Our previous definition of *ϵ* ^2^(*x*) (Eq. 6) represents the positional error from a single independent measurement of the morphogen concentration at a specific location in space. However, organisms may further improve the positional information of the morphogen gradient through space-time averaging.

To incorporate the scenario of space-time averaging, we assume that the cell at position *x* uses a sensor with size *a* to detect the morphogen concentration over the space interval (*x* −*a, x* + *a*) for time *T* (Fig. 3a). Then the space-time averaged molecular count is written as

**FIG. 3.**
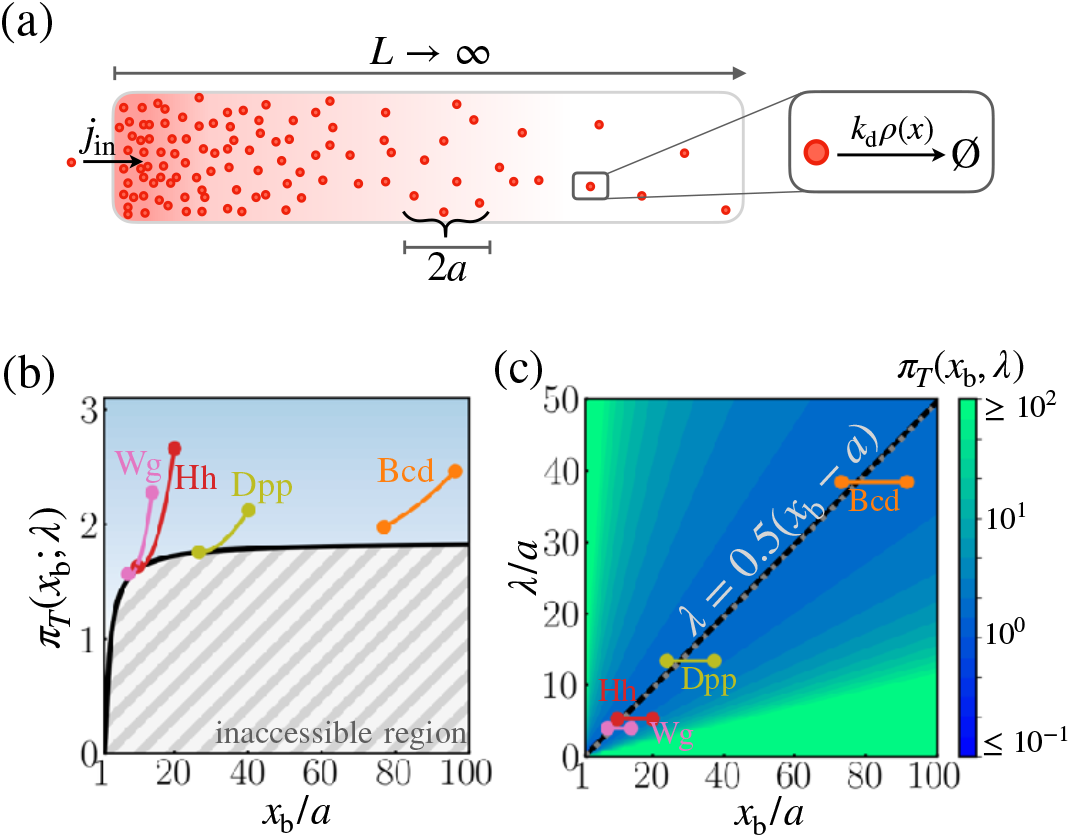
Cost-precision trade-off with the space-time averaged measurement. **(a)** Schematic of the model. **(b)** The optimal trade-off product *π*_*T*_ (*x*_b_) obtained numerically with respect to the location of the target boundary position (*x*_b_*/a*). The grey hashed regions represent the inaccessible regions. Shown are the trade-off products of the morphogens profiles of Bcd, Wg, Hh, and Dpp. **(c)** The black line denotes the optimal characteristic decay length (λ_min_) with respect to the position *x*_b_*/a*. The color scale indicates the trade-off product *π*_*T*_ (*x*_b_) computed for each pair of λ*/a* and *x*_b_*/a* values. In **b** and **c**, the depicted trade-off product is normalized by *α*_*o*_. The grey dotted line depicts the linear approximation of λ_min_. The parameters for the naturally occurring morphogen profiles are further described in the SI Appendix **Length scales of Bcd, Wg, Hh, and Dpp**.

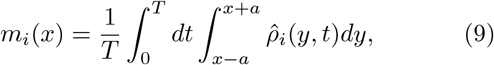

where 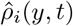 represents an estimate of molecular count at position *y* from an *i*-th independent measurement. Repeated measurements (*N* ≫ 1) leads to the mean value of molecular counts

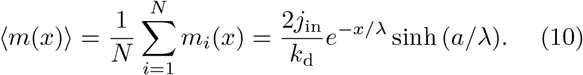

In this case, *j*_in_ and *k*_d_ are in the units of #*/*time and time^−1^, respectively. With a sufficiently long measurement time 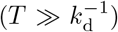, the variance of *m*(*x*) can be approximated to^19^

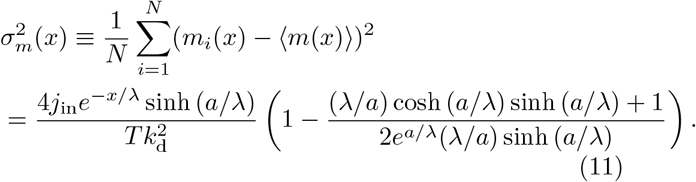

Analogously to Eq. 6, the squared relative error (precision) at position *x* can be quantified as follows,

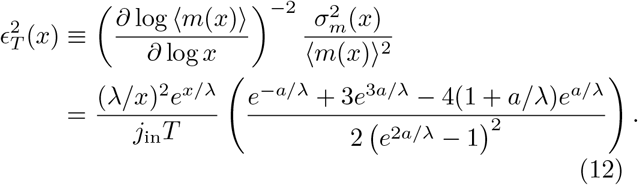

The cost of maintaining the morphogen profile at steady states is proportional to *j*_in_*T*, which simply represents the total number of morphogens produced for *T*, such that the total thermodynamic cost for producing morphogens for time *T* is

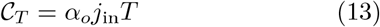

In the product between the net morphogen production and the positional error from the averaged signal, the number of morphogens produced for time *T, j*_in_*T* cancels off, which yields the following expression of the cost-precision trade-off:

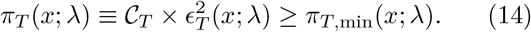

With a given target boundary = and the sensor size *a*, the value of *π*_*T*_ (*x*_b_; λ) is solely determined by λ, analogously to *π*_*o*_(*x*_b_; λ). Thus, the lower bound of *π*_*T*_ (*x*_b_), i.e., *π* _*T*,min_ (*x*_b_), can be determined simply by tuning λ to a value that minimizes *π*_*T*_ (*x*_b_; λ), i.e., λ_min_(*x*_b_). Both *π*_*T*,min_(*x*_b_) and λ_min_(*x*_b_) increase monotonically with *x*_b_. In particular, λ_min_(*x*_b_) is constrained by the inequalities (*x*_b_ — *a*)*/*2 λ_min_(*x*_b_) ≤ *x*_b_*/*2 (see Eqs. S31 and S32 for derivation). The boundary of the inaccessible region in Fig. 3b constrains *π*_*T*_ (*x*_b_), suggesting that there is a minimal morphogen production for a given positional error. For instance, in order to suppress the positional error down to 10 % (*ϵ*_*T*_ (*x*_b_) ≈ 0.1) at *x*_b_ = 60*a*, the characteristic length scale must be ∼30*a*, which demands that the system synthesize at least ∼182 morphogen molecules (Fig. 3b).

As we have done for *π*_*o*_, we can evaluate *π*_*T*_ of the four morphogen profiles of Bcd, Wg, Hh, and Dpp at their target boundaries (*x*_b_), by using the cell size as proxies for the sensor size, and by assuming that the morphogen profile is measured for a sufficiently long time (i.e.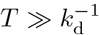, the validity of which is further discussed in the SI Appendix **Time scales of Bcd, Wg, Hh, and Dpp**). We find that the characteristic decay lengths 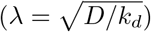 of all four morphogen profiles are close to their respective λ_min_(*x*_b_) values (Fig. 3c). Thus, from the space-time averaged measurement of morphogen profiles, the four morphogen profiles are also formed with nearly optimal cost-precision trade-off condition (Fig. 3b).

## DISCUSSION

### Comparison of the trade-off products, *π*_*o*_ and *π*_*T*_

Whereas the two models of biological pattern formation seem to differ in the definitions of both the precision and the associated cost, the measure of precision in the two models are in fact limiting expressions of each other. The concentration detected in the measurement with space-time averaging, defined as *m*(*x*)*/*(2*a*) at the limit of small sensor size (*a/*λ ≪ 1) (Eq. 10) is identical to the one in the point measurement in the limit of *L* ≫ λ (Eq. S2), both yielding 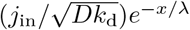. On the one hand, *ϵ* ^2^(*x*), defined through *ρ*_ss_(*x*), is experimentally quantifiable through repeated measurements of the morphogen profile from fixed images^4,5,49^, On the other hand, 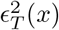, defined through *m*(*x*), may represent the precision accessible to cells that detect morphogens through receptors on the cell surface.

The cost associated with the morphogen profile in the case of point measurement, 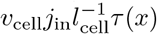, is a quantity that reflects the morphogen production required to reach steady state at position *x*. In contrast, the cost of morphogen production with space-time averaging, *j*_in_*T*, integrated over a time interval 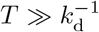, is identical for all positions. The latter assumption is required in the derivation of the expression of 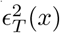 (see Appendix 1 of ref.^19^). We additionally demand *T* ≫ *τ* (*x*) in order for *j*_in_ *T* to represent the total cost of morphogen profile formation and maintenance. However, *T* may not necessarily be larger than either 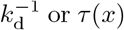. For instance, the Bcd profile is degraded and established at time scales of ∼ 1 hour^13,50^, but must be measured within 10 minutes^51^ (see further discussion in the SI Appendix **Time scales of Bcd, Wg, Hh, and Dpp**). The energetic cost of naturally occurring morphogen profiles is likely determined in between the two limiting cases.

*π*_*o*_(*x*) and *π*_*T*_ (*x*) represent cost-precision trade-off relations acquired from two limiting scenarios of morphogen detection. For actual biological systems, cost-precision trade-off involving the pattern formation is presumably at work in between the two scenarios. The finding that the two limiting models can be effectively minimized by similar λ values at a given position is of great significance.

### The entropic cost of forming precise morphogen profiles

Biological pattern formation by morphogen gradients is a process operating out of equilibrium^20^, in which thermodynamic cost is incurred to produce a morphogen gradient with minimal error for the transfer of positional information against stochastic fluctuations. In the SI Appendix **Reversible reaction-diffusion model of morphogen dynamics**, we extend our discussion of cost-precision trade-off on the models that include unidirectional irreversible steps in the synthesis and degradation by considering a more general bi-directional reversible reaction kinetics where the forward and reverse rates are well defined at every elementary process, which allows us to quantify the entropy production rate (thermodynamic cost) for the formation of morphogen gradient,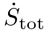. Similar to the point measurement, we derive expressions for the relaxation time to the steady state (*τ*_rev_(*x*)), the positional error 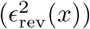, and the mean entropy production rate 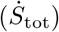. The trade-off among the thermodynamic cost, speed of formation 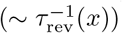, and precision is quantified by the product,

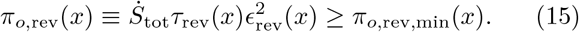

At a boundary *x* = *x*_b_, the lower bound *π*_*o*,rev,min_ (*x*_*b*_), increases monotonically with *x*_b_. With 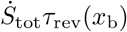 representing the entropy production during the time scale associated with the morphogen profile formation, Eq. 15 is reminiscent of the thermodynamic uncertainty relation (TUR), a fundamental trade-off relation between the entropy production and the precision of a current-like output observable for dynamical processes generated in nonequilibrium with a universal bound^52–58^. However, unlike TUR which has a model independent lower bound, the lower bound of Eq. 15 is model-specific and position dependent. The cost-precision trade-off of pattern formation discussed in this study fundamentally differs from that of TUR^58–61^, in that the concentration profile of morphogen is not a current-like observable with an odd-parity with time-reversal; however, still of great significance is the discovery of the underlying principle that the pattern formation is quantitatively bounded by the dissipation. For future work, it would be of great interest to consider employing the morphogen-induced currents through signaling pathways, such as transcription, as the output observable to which TUR can be directly applied.

### Thermodynamic cost is a key physical constraint for morphogen profile formation

There are multiple possibilities of achieving the same critical threshold concentration *ρ*(*x*_b_) at the target boundary *x* = *x*_b_ through the combination of morphogen synthesis rate (*j*_in_), diffusivity (*D*) and degradation rate (*k*_*d*_) (inset in Fig. 4). A steeper morphogen profile with larger *j*_in_ and smaller 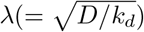 (red in Fig. 4) leads to a more precise boundary but it incurs a higher thermodynamic cost. The opposite case with a morphogen profile with smaller *j*_in_ and larger λ gives rise to a less precise boundary with a lower cost (brown in Fig. 4). As the main result, we show that λ can be tuned to minimize the cost-precision trade-off product, which leads to the formation of cost-effective morphogen profiles. Specifically, λ ≈ (0.43 − 0.50)*x*_b_, which suggests that the target boundary of the exponentially decaying morphogen profile, *c*(*x*) = *c*_0_*e*^−*x/*λ^, should be formed at *c*(*x*_b_) ≈ 0.1*c*_0_. Remarkably, the λ’s of naturally occurring morphogen profiles in fruit fly development are close to their respective optimal values, and we confirm that the target boundaries of biological pattern from those profiles are identified at *c*(*x*_b_) ≈ 0.1*c*_0_ (see Fig. S4). These findings lend support to the hypothesis that along with the reduction in the positional error, the thermodynamic cost is a key physical constraint in designing the morphogen profiles.

**FIG. 4.**
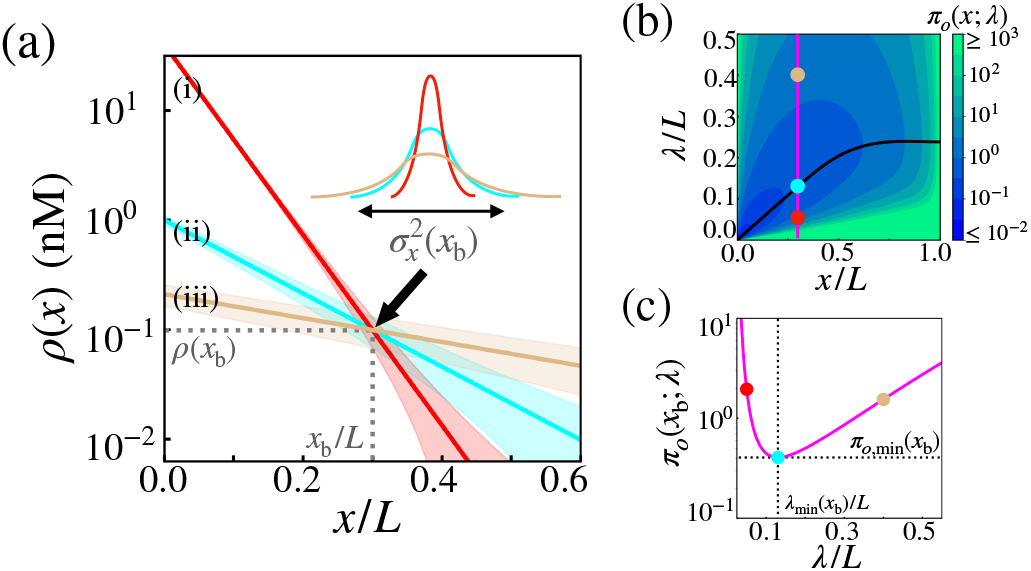
Optimal concentration profile of morphogens. **(a)** Three possible morphogen profiles (i) (red), (ii) (cyan), (iii) (brown) with different λ’s (λ^(i)^ > λ^(ii)^ > λ^(iii)^), generated with different values of morphogen influxes 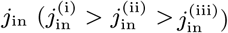. The morphogen concentration of the three profiles coincide at *x*_b_, giving rise to the same threshold value *ρ*(*x*_b_) but different positional errors (*ϵ*^2^(*x*_b_; λ)). **(b)** The diagram of trade-off product for the point measurement, *π*_*o*_(*x*; λ), plotted with respect to *x* and λ. The black line indicates the optimal decay length, λ_min_ at position *x*. Shown on the diagram are the trade-off product *π*_*o*_’s for the three cases. **(c)** The value of *π*_*o*_ as a function of λ at *x* = *x*_b_. The trade-off product is minimized to 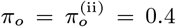 with 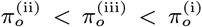. The morphogen profile of the case (ii) displays the lowest positional error for a given thermodynamic cost.

### Concluding remarks

The classical SDD model offers a simple and powerful framework to study basic properties of morphogen dynamics^4–6,11,13,19,25^. The molecular mechanisms underlying the formation of morphogen profiles are, however, much more complex than those discussed here. Even the seemingly simple diffusive spreading of the morphogen can originate from many different mechanisms^62^. In fact, among the four biological examples shown in Figs. 2 and 3, it has been suggested that Wg and Hh spread over the space through active transport mechanisms rather than through passive diffusion^19,63–65^. Furthermore, it is known that multiple morphogen profiles relay combinatorial input signals that determine the expressions of an array of downstream target genes and their subsequent interactions^66–68^. Ongoing studies employing model systems including the fruit fly, zebrafish, chicken, and mouse are providing key insights into complex tissue patterning mechanisms^9,10,69–73^.

Modified trade-off relations in systems with more complex geometries, such as the 1D model with a distributed morphogen source, and the morphogen dynamics on a sphere can be conceived as well (see SI Appendix). For the latter case, the λ values, for the morphogens inducing the endoderm and mesoderm of zebrafish embryos, are found far greater than those leading to the trade-off bound (Fig. S3b). The large trade-off product values may either simply indicate that the thermodynamic cost is not necessarily a key physical constraint or reflect the presence of other molecular players in a more complex mechanism establishing the zebrafish germ layers.^69,74–76^. For future work, it would be of great interest to assess the cost-effectiveness of more complex systems through the theoretical framework of cost-precision trade-off proposed in this study.

## Acknowledgments

This work was supported by the KIAS individual Grants CG067102 (Y.S.) and CG035003 (C.H.) at Korea Institute for Advanced Study. We thank the Center for Advanced Computation in KIAS for providing computing resources.

## SUPPLEMENTARY INFORMATION

### 1. LOCALIZED SYNTHESIS-DIFFUSION-DEPLETION MODEL IN A ONE-DIMENSIONAL ARRAY

Here we provide detailed derivations of the cost-precision trade-off relation for the localized synthesis-diffusion-depletion model of the morphogen dynamics in a one-dimensional array of cells. We begin with the reversible reaction-diffusion dynamics of morphogen profile formation, where the forward and reverse rates are well defined at every elementary step, which enables the calculation of the entropy production rate per cell 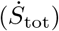. Through the discussion of the reversible process, we provide mathematical details on the derivation of the cost-precision trade-off relation for the irreversible reaction diffusion process of morphogen synthesis and depletion presented in the main text. Next, we derive the conditions that correspond to the minimum trade-off product for the irreversible process with point (*π*_*o*_) and space-time averaged (*π*_*T*_) measurements.

#### A. Reversible reaction-diffusion model of morphogen dynamics

We explore the trade-offs among the speed, precision, and entropy production in the reversible morphogen dynamics. Similarly to the irreversible morphogen dynamics presented in the main text, the dynamics is defined on a one-dimensional array of cells. We denote the amount of morphogen in the cell of the volume (*v*_cell_) and size (*l*_cell_) at the interval between *x* and *x* + *l*_cell_ by the concentration, *ρ*(*x, t*) [conc ≡#*/v*_cell_]. At the left boundary (see Fig.S1a), the morphogen is injected with rate *j*_in_ [conc × *l*_cell_*/*time], and taken out by the corresponding reverse reaction with rate *v*_out_ [*l*_cell_*/*time]. At all positions, the morphogen is depleted with rate *k*_d_ [time^−1^], and synthesized by the corresponding reverse reaction with rate *r*_s_ [conc*/*time]. Additionally, the morphogen spreads across space with diffusivity 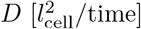. At the continuum limit (i.e. with *l*_cell_ ≪ *L*), the dynamics of the morphogen is described by the partial differential equation

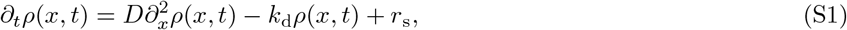

with two boundary conditions at *x* = 0 and *x* = *L*, −*D∂_*x*_ρ*(*x, t*)|_*x*=0_ = *j*_*in*_ −*v*_*out*_ *ρ*(0, *t*) and −*D∂_*x*_ρ*(*x, t*))|_*x*=*L*_ = 0. The characteristic length scale of the dynamics associated with Eq. S1 is set by 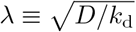. Then, the steady state concentration can be written as

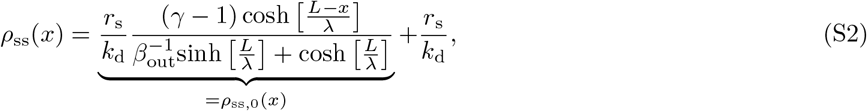

where *ρ*_ss,0_(*x*) denotes the position-dependent portion of the concentration profile. We introduce three dimensionless parameters: *β*_in_ ≡ *j*_in_*/*(λ*r*_s_), *β*_out_ ≡ *v*_out_*/*(λ*k*_d_), and their ratio *γ* ≡ *β*_in_*/β*_out_ which is related with the thermodynamic cost (see Eq. S20). We only consider the regime with *γ >* 1, where a net positive current of morphogen is supplied at the left boundary. In what follows, we will derive the quantitative expressions of the (i) *speed*, (ii) *precision*, and (iii) *thermodynamic cost* associated with the morphogen profile.

##### 1. Speed

The concentration profile varies with time to reach its steady state, *ρ*_ss_(*x*) ≡lim_*t →∞*_ *ρ*(*x, t*), from the initial profile *ρ*(*x*, 0) = 0. The time evolution of *ρ*(*x, t*) can be analyzed by solving the Eq. S1 in Laplace domain:

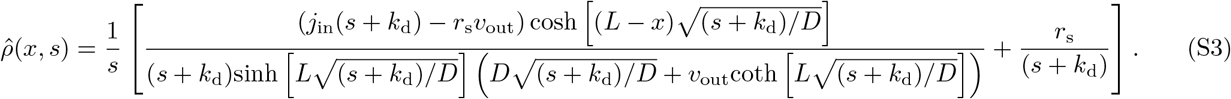

The expression of the steady state profile (Eq. S2) is obtained using

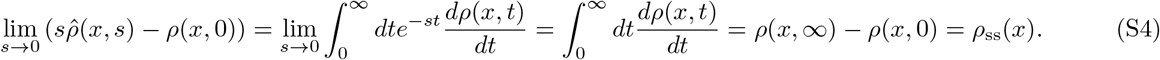

To characterize the relaxation dynamics to the steady state profile, we define the relaxation function

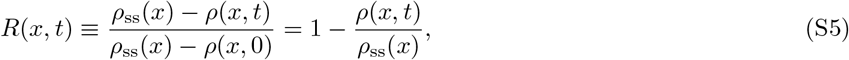

which monotonically decays from 1 to 0 at all positions as *t*→ ∞. The associated mean relaxation time at position *x, τ*_rev_(*x*), is given by

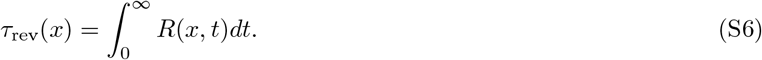

Since the Laplace transform of the relaxation function is 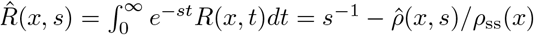, the local accumulation time at position *x*, 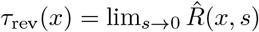, is obtained as

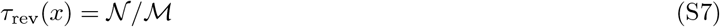

with

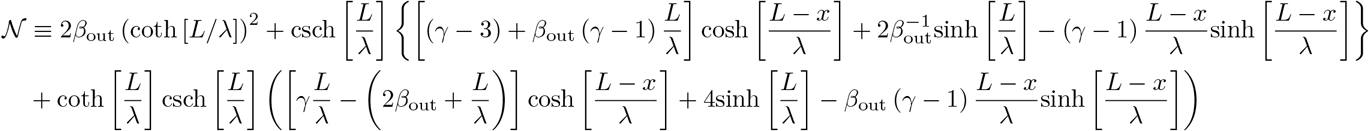

and

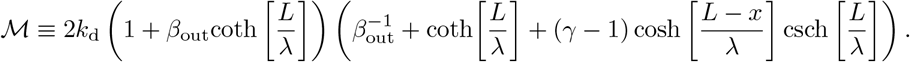

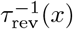 quantifies the speed at which the morphogen profile is established at position *x*^13,77^. *The expression for the characteristic time in the irreversible morphogen dynamics is obtained when r*_s_ and *v*_out_ are negligibly small. At the limit of *r*_s_ → 0 and *v*_out_ → 0, with the substitutions *β*_out_ = *v*_out_*/*(λ*k*_d_) and *γ* = *j*_in_*k*_d_*/*(*v*_out_*r*_s_), Eq. S7 is simplified to

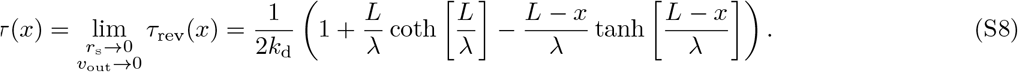

##### 2. Precision

The local measure of precision at *x* is defined as the squared relative positional error (Eq. 1),

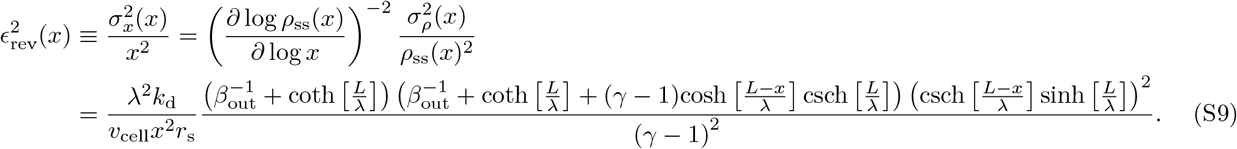

The expression for the squared relative error in the irreversible morphogen dynamics can be obtained at the limit of *r*_s_ → 0 and *v*_out_ → 0, with the substitutions *β*_out_ = *v*_out_*/*(λ*k*_d_) and *γ* = *j*_in_*k*_d_*/*(*v*_out_*r*_s_):

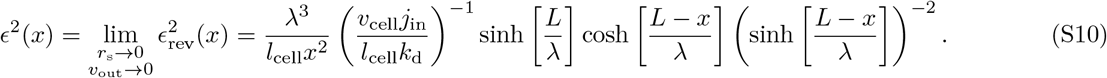

##### 3. Thermodynamic cost

The total entropy production rate required to maintain the steady state morphogen profile is given by

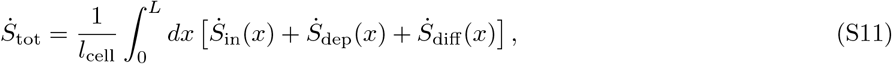

where the three components of the position-dependent entropy production rate are

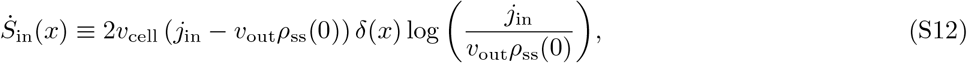

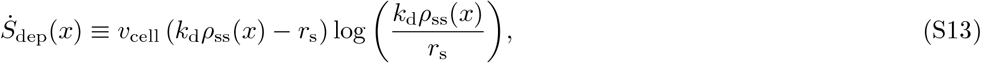

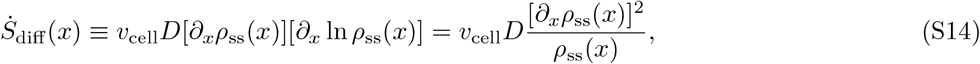

with the Boltzmann constant set to unity (*k*_*B*_ = 1). Eqs. S12 and S13 represent the entropy production rates from the net morphogen current flowing in and out of the system, respectively, in the Schnakenberg formalism. Similarly, Eq. S14 arrises from the diffusive flux^20^. Briefly, Eq. S14 can be understood as 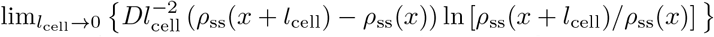, in which 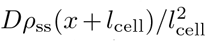 and 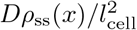 represent the forward and reverse currents between adjacent spatial compartments in the discretized spatial representation. It is noteworthy that the asymmetry of concentrations generates a flow of morphogen current between neighboring cells, and contributes to the entropy production. If the morphogen profile is uniform in space, which occurs at the detailed balance condition 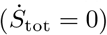, the cells cannot infer any spatial information from the morphogen.

To obtain a simplified expression of 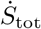, we use the following two equalities. First, at steady state, the net injection of the morphogen at the left-boundary must be equal to the net depletion of the morphogen occurring at all positions, so that

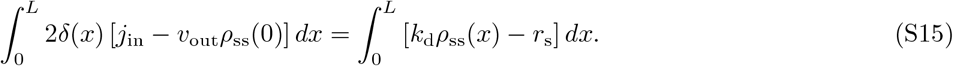

Second, 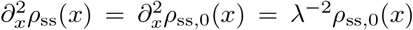, with *ρ*_ss,0_ defined in Eq. S2, lead to the following equality

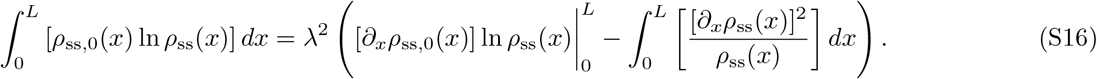

Then, 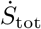 can be expressed as follows:

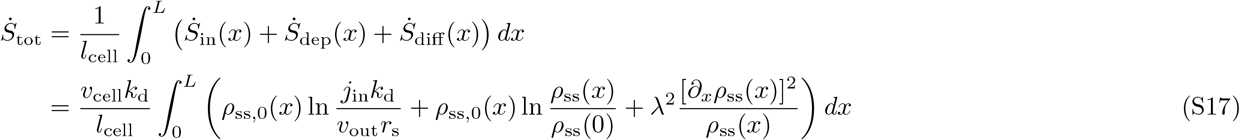

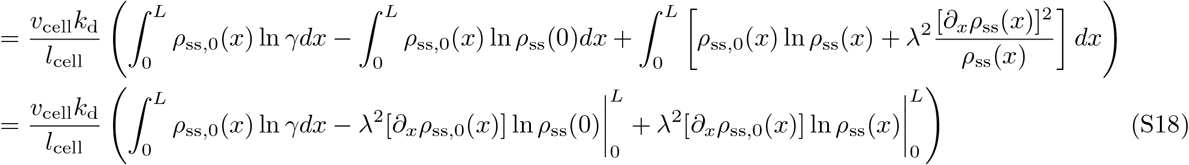

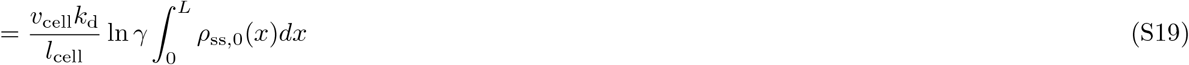

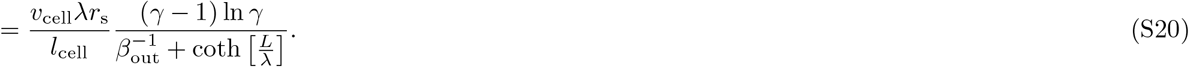

We used S15 and S16 to obtain S17 and S18, respectively. Next, the boundary condition of the system, *∂*_*x*_*ρ*(*x*)|_*x*=*L*_ = 0, leads to S19. Finally, the evaluating the integral of *ρ*_ss,0_(*x*) leads to the last expression.

In the irreversible system with negligibly small *r*_s_ and *v*_out_, the entropy production rate is undefined, since 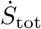 (S20) diverges with ln *γ* → ∞. Nevertheless, the mean morphogen production rate is still related with 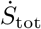 normalized by ln *γ* at the limit of *r*_s_ → 0 and *v*_out_ → 0:

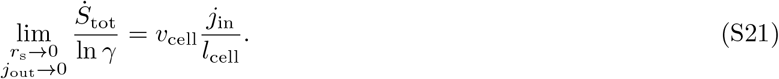

**FIG. S1.**
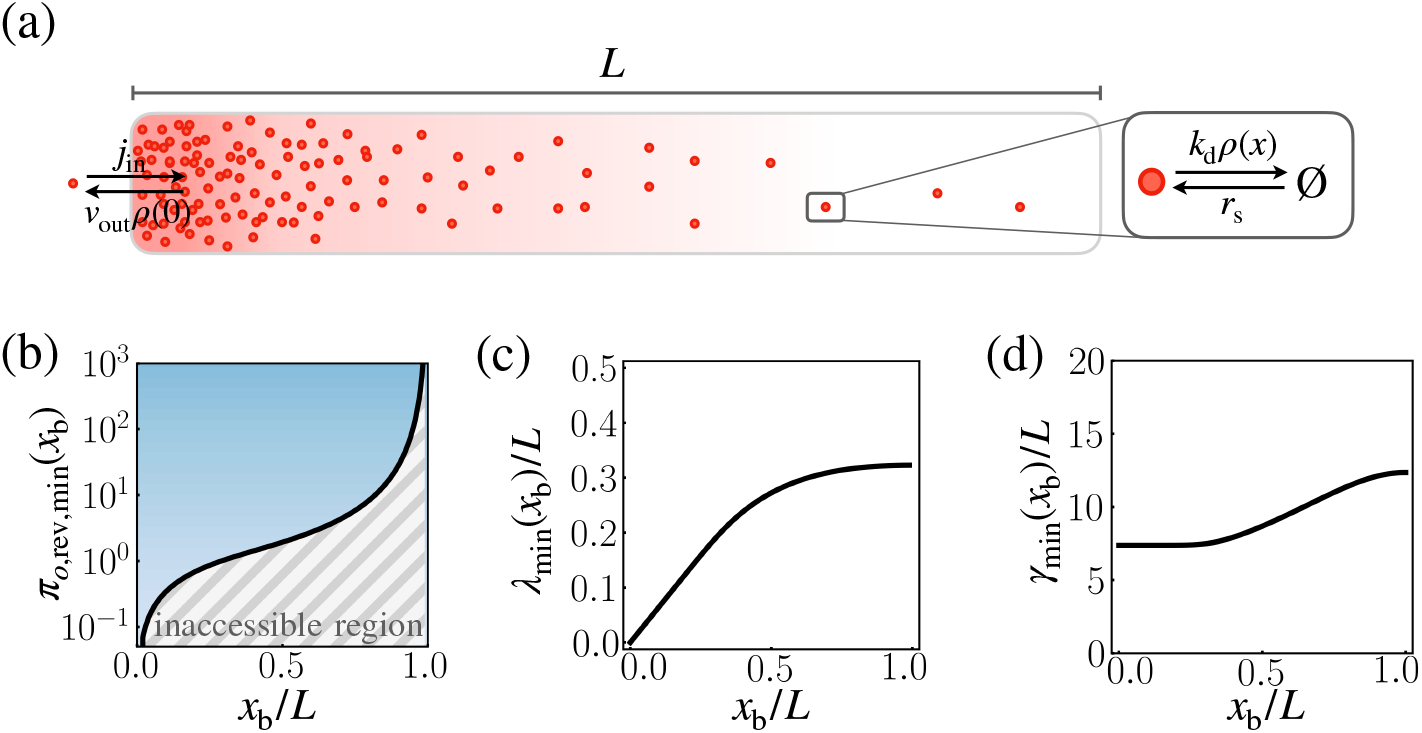
Speed-precision-cost trade-off for the reversible morphogen dynamics. **(a)** Schematic of the reversible reaction-diffusion dynamics of morphogen profile formation. **(b)** The solid black line depicts the optimal trade-off product *π*_*o*,rev,min_(*x*_b_), obtained numerically by minimizing with respect to λ and *γ*, with *β*_out_ → ∞. The trade-off product is normalized by *L/l*_cell_. **(c)(d)** The optimal characteristic decay length λ_min_ (b) and thermodynamic drive *γ*_min_ (c) corresponding to *π*_*o*,rev,min_(*x*_b_).

##### 4. Trade-off product

The product of the local accumulation time (*τ*_rev_(*x*), Eq. S7), the squared relative positional error (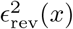, Eq. S9), and the mean entropy production rate per cell (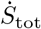, Eq. S20) leads to the position-dependent trade-off product *π*_*o*,rev_(*x*),

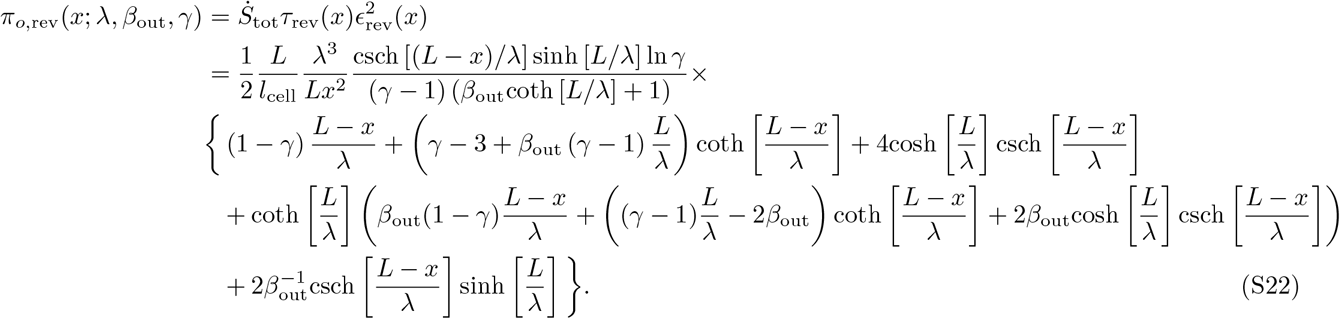

With fixed *L, l*_cell_, and *x* = *x*_b_, the lower bound of *π*_*o*,rev_(*x*_b_; λ, *β*_out_, *γ*), *π*_rev,min_(*x*_b_), can be obtained by minimizing *π*_*o*,rev_(*x*_b_; λ, *β*_out_, *γ*) with respect to *γ, β*_out_, and λ. For a fixed value of *γ >* 1, the partial derivative of *π*_*o*,rev_(*x*_b_; λ, *β*_out_, *γ*) with respect to *β*_out_ is always smaller than 0,

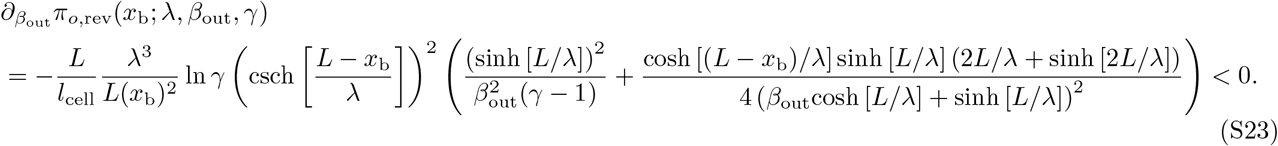

Thus, *π*_*o*,rev_(*x*_b_) is minimized at the limit of large *β*_out_ (or, equivalently, large *β*_in_ = *γβ*_out_), with the following expression:

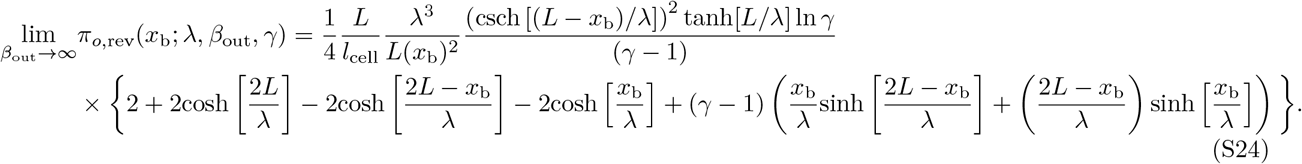

With the characteristic decay length set to λ = *x*_b_, it can be shown that 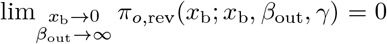. Since *π*_*o*,rev,min_(*x*_b_) ≤ *π*_*o*,rev_(*x*_b_; *x*_b_, *β*_out_, *γ*) by the definition of λ_min_(*x*_b_), *π*_*o*,rev,min_(0) = 0.

For *x*_b_ > 0, the optimal trade-off product, *π*_rev,min_(*x*_b_), and the corresponding values of λ_min_(*x*_b_) and *γ*_min_(*x*_b_), were numerically obtained (Fig. S1). *π*_*o*,rev,min_(*x*_b_) constrains dynamical properties of the morphogen profile, specifying, for instance, the minimum entropy production rate for a given speed and precision (Fig. S1b). With a small *π*_*o*,rev_(*x*_b_; λ, *β*_out_, *γ*) close to *π*_*o*,rev,min_(*x*_b_), the system can rapidly generate precise morphogen profiles at *x*_b_ with low thermodynamic cost. In general, *π*_*o*,rev,min_(*x*_b_), λ_min_(*x*_b_) and *γ*_min_(*x*_b_) all increase monotonically with *x*_b_ (Fig. S1b,c,d).

#### B. Minimum trade-off product of the point measurement for the irreversible reaction-diffusion process

We provide the mathematical details involving the trade-off product of the point measurement (*π*_*o*_, Eq. 7 and Fig. 2 in the main text). With the expressions derived in the SI Appendix **Reversible reaction-diffusion model of morphogen dynamics**, the thermodynamic cost of morphogen production required to reach the steady state concentration at position *x* (Eq.5 in the main text) amounts to the product between Eq. S8 and Eq. S21,

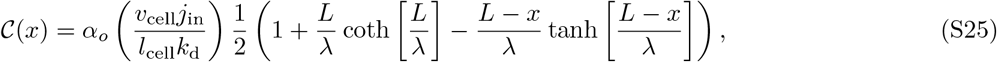

and the squared relative positional error, which is identical to Eq. S10, is

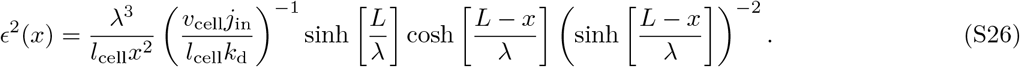

Then, the trade-off product reads

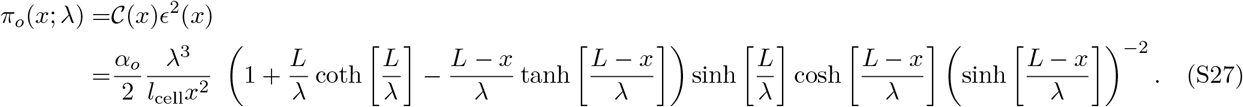

For fixed *x* = *x*_b_, *π*_*o*_(*x*_b_; λ) is numerically minimized with respect to λ to give rise to the solid black line in Fig. 2c. When *L/*λ ≫ 1, the trade-off product (Eq. S27) is approximated to

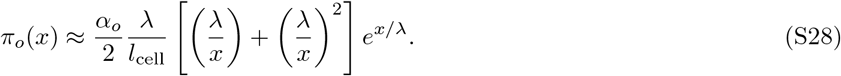

For a given position 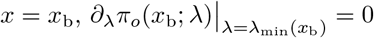 determines 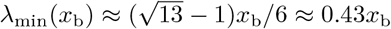 (grey dotted line in Fig. 2c).

#### C. Minimum trade-off product of the space-time-averaged measurement for the irreversible reaction-diffusion process

As in the main text, the trade-off product from the space-time averaged measurement is given by

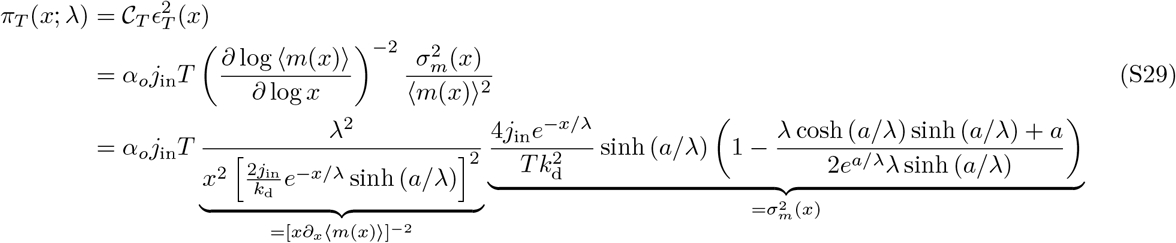

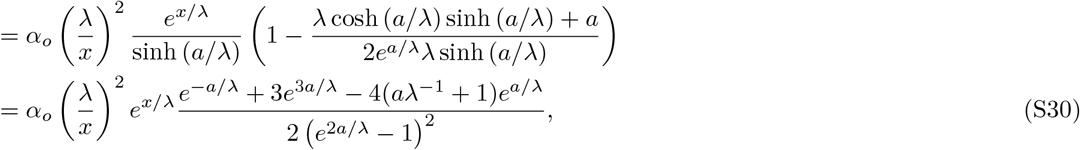

The *π*_*T*_ (*x*; λ) minimizing λ at *x* = *x*_b_, namely λ_min_(*x*_b_), is found at the interval (*x*_b_ − *a*)*/*2 ≤ λ_min_(*x*_b_) ≤ *x*_b_*/*2, since

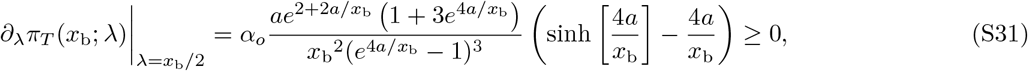

and

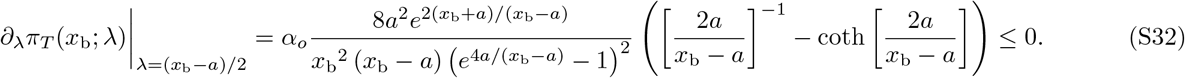

For Fig. 3 in the main text, the exact values of *π*_*T*,min_(*x*_b_) and λ_min_(*x*_b_) were obtained by numerically minimizing Eq. S30 with respect to λ.

### II. DISTRIBUTED MORPHOGEN SOURCE

We consider a case where the source of morphogen production is distributed over the space in the irreversible reaction-diffusion model of morphogen dynamics. The space-time evolution of the morphogen is described by

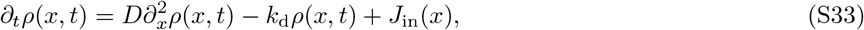

with the boundary conditions at *∂*_*x*_*ρ*(*x, t*)| _*x*=0,*L*_ = 0. The production term, *J*_in_(*x*) is assumed to be exponentially distributed with a characteristic length λ_p_,

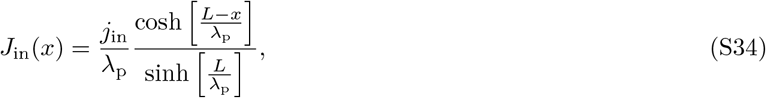

so that 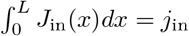. The solution to Eq. S33 in the Laplace domain is given by

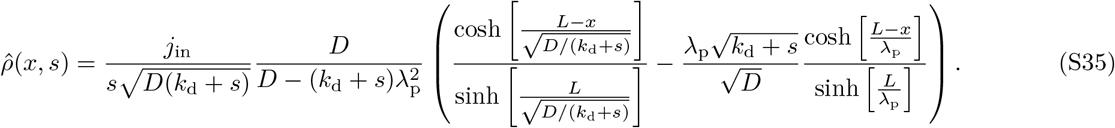

By following a procedure similar to that shown in the section **Reversible reaction-diffusion model of morphogen dynamics**, we can obtain the following expressions for the steady state profile (*ρ*_ss_(*x*), Eq. S4), and the precision (*ϵ*^2^(*x*), Eq. S9):

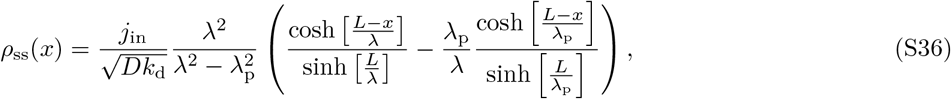

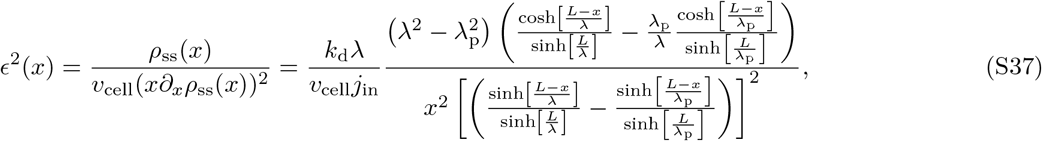

with 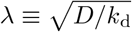. Next, the characteristic timescale to approach the steady state profile (*τ* (*x*), Eq. S7) can be written as

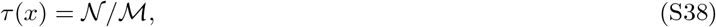

with

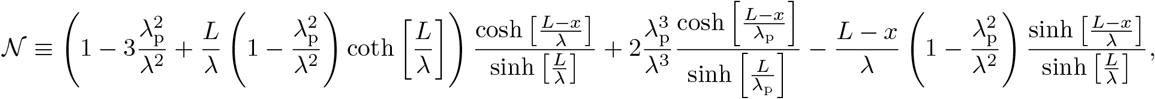

and

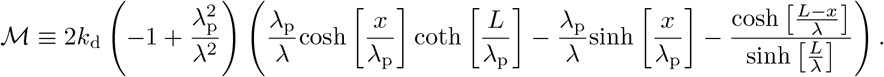

Then, the trade-off product is expressed as

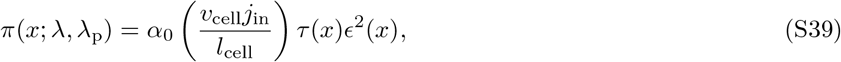

with the proportionality constant *α*_0_. For fixed *x* = *x*_b_, *L*, and λ_p_, the trade-off product can be minimized by tuning λ to the optimal value, λ_min_(*x*_b_). As λ_p_ increases, the optimal product and the associated optimal characteristic length λ_min_(*x*) decrease (Fig. S2). In Fig. S2, the characteristic length scale λ_d_ is defined as 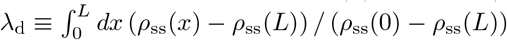.

### III. DYNAMICS ON A SPHERE

We consider the morphogen profile formation on the surface of a sphere, the situation of which the dynamics of Cyclops, Squint, and Fgf8 in the developing zebrafish embryo at ∼50% epiboly are relevant. When the concentration of the morphogen in a cell with spherical coordinates (*r, θ, ϕ*) is denoted by *ρ*(*r, θ, ϕ*), with the symmetry along the azimuthal angle and the condition of the morphogen dynamics being confined to the spherical shell of radius *r* = *R* (i.e. *∂*_*ϕ*_*ρ* = 0, *∂*_*r*_*ρ* = 0, and *r* = *R*) the morphogen concentration satisfies the following evolution equation:

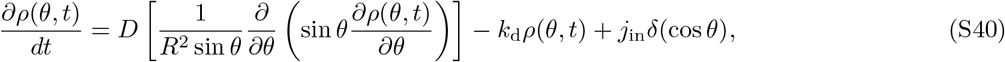

with boundary conditions at *∂*_*θ*_*ρ*|_*θ*=0,*π*_ = 0. The morphogen dynamics is fully specified by the system size *R* [*l*_*cell*_], the depletion rate *k*_d_ [1*/*time], the diffusion coefficient 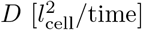, and the injection rate *j*_in_ [conc*/*time]. The last term represents the influx of morphogens due to their synthesis by the “band” of cells at the equator (Fig. S3a). With the substitution *z* = cos *θ* and the initial condition *ρ*(*z, t*) = 0, the solution of Eq. S40 in the Laplace domain with 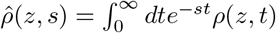, can be obtained by solving

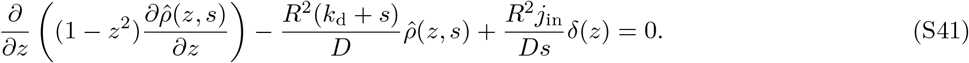

The solution to the differential equation of this type can be expressed as 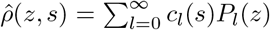, where *P*_*l*_(*z*) (*l* = 0, 1, …) is the Legendre polynomial, satisfying 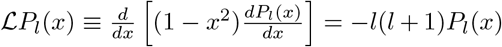, with the orthogonality condition 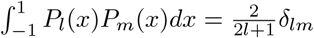. To find the coefficient *c*_*l*_(*s*), we rearrange Eq. S41 into

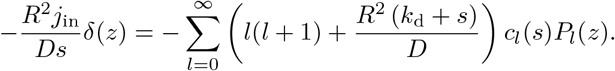

**FIG. S2.**
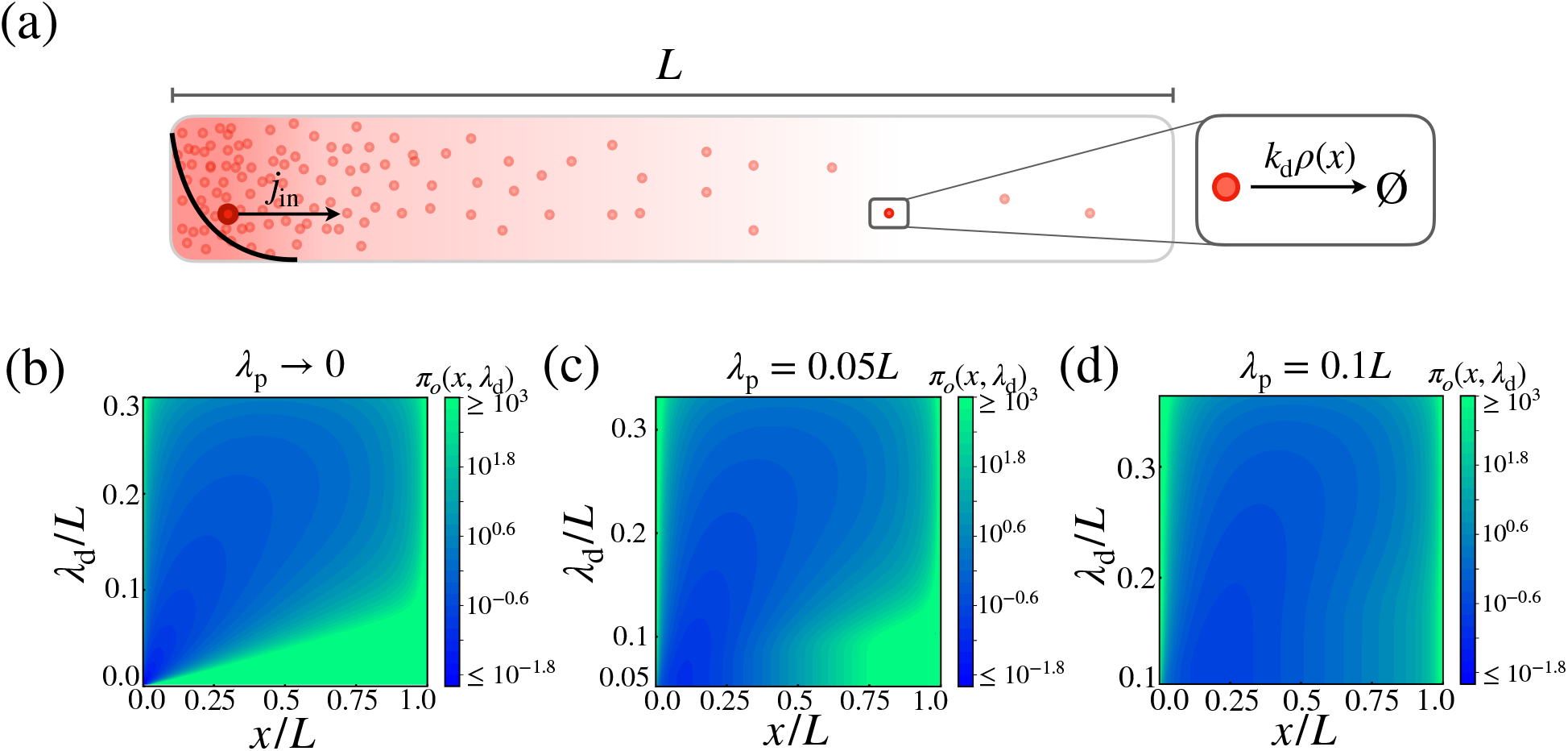
Cost-precision trade-off in the morphogen dynamics with distributed synthesis. **(a)** Schematic of the distributed synthesis of the morphogen. The black solid line represents the morphogen synthesis rate which is roughly exponentially distributed with characteristic decay length λ_p_. **(b)(c)(d)** The trade-off values (*π*_*o*_(*x*, λ_d_)) for (b) λ_p_ = 0, (c) λ_p_ = 0.05*L*, and (d) λ_p_ = 0.1*L*. The color scale indicates the trade-off product *π*_*o*_ normalized by *α*_*o*_*L/l*_cell_. The formula for the *π*_*o*_ values are given in SI Appendix **Distributed morphogen source**. The characteristic length scale is defined as 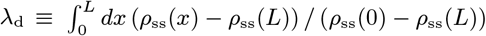.

By using the orthogonality condition of Legendre polynomials, we obtain the solution of Eq. S41,

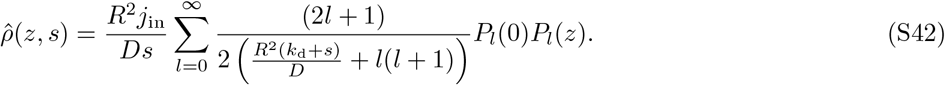

The steady state concentration can be expressed as 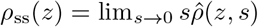.

The average relaxation time can be calculated using 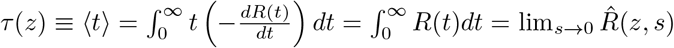, which yields

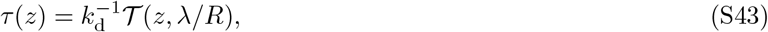

with 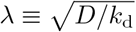 and

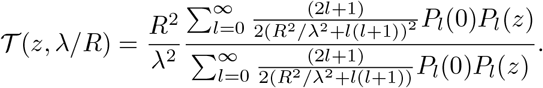

The number of morphogens produced from the cells at the equator per unit time is *v*_cell_(2*πR/l*_cell_)*j*_in_. Thus, the thermodynamic cost to approach the steady state profile at *z* is

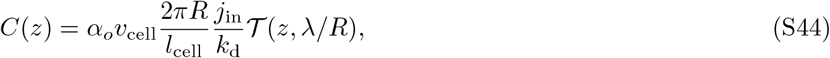

where *α*_*o*_ is a proportionality constant.

Next, using 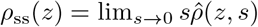 (Eq. S42), one gets the local precision of the morphogen profile at position *z*,

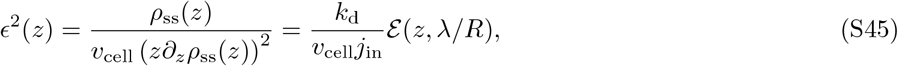

**FIG. S3.**
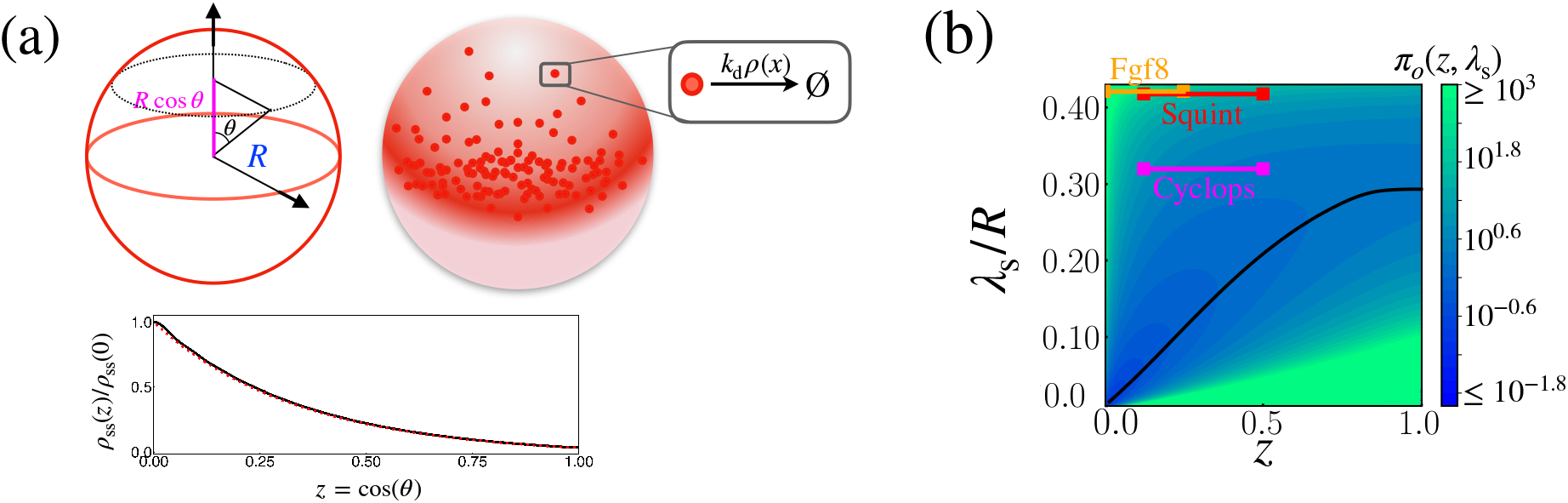
Cost-precision trade-off of the morphogen dynamics on a sphere. **(a)** (Top) Schematic of the morphogen synthesized from the equator of the spherical shell (red region). (Bottom) The relative steady state concentration profile *ρ*_ss_(*z*) when *R*^2^*/*λ^2^ = 10. The solid black line was computed from evaluating 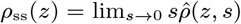 (Eq. S42) up to *l* = 84. The dotted red line was obtained by numerically solving Eq. S40. **(b)** The trade-off product (*π*_*o*_(*z*, λ_s_)) for pairs of target position *z* and characteristic length scale λ_s_ values, normalized by *α*_*o*_(2*πR/l*_cell_). The formula for *π*_*o*_ is given in SI Appendix **Dynamics dynamics on a sphere**. The characteristic length scale λ_s_ is defined as 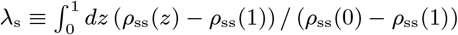.

where

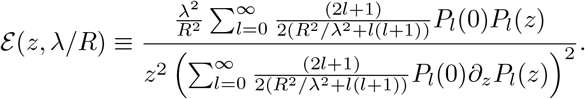

Finally, the trade-off product, which is plotted in Fig. S3b as a function of *z* and λ_*s*_, is defined as

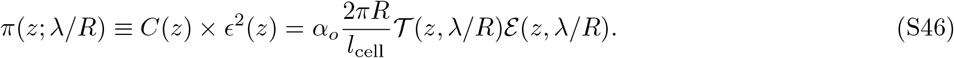

The length scale λ_s_ used in the plot is defined as 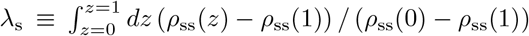. At the targetposition *z* = *z*_b_, the trade-off product can be minimized by tuning λ to the optimal value λ_min_, which is plotted in black in Fig. S3b.

### IV. LENGTH SCALES OF BCD, WG, HH, AND DPP

Here we explain various length scales of naturally occurring morphogen profiles of *Drosophila* (Fig. S4 and Table. S1).

- **Bcd**. The characteristic decay length of Bcd was set to λ_Bcd_ = 100 *µ*m as reported in^4^. The system length *L* = 475 *µ*m and sensor radius *a* = 2.6 *µ*m was obtained from Fig. 3A of Ref.^4^. The boundary position of the Bcd target *hb* expression was set to *x*_b_ = 190 ∼ 238 *µ*m (Fig. 3C of Ref.^42^).
- **Wg**. The characteristic decay length of Wg was set to λ_Wg_ = 6 *µ*m (Ref.^5^). Although the Wg dynamics reported in Fig. S4b is not from the endogenous concentration profile, we assume that the wild type profile forms with similar length and time scales. While Wg induces the expression of multiple genes, including *distalless* and *vestigial*, we picked *sens* for our analysis because its expression profile was quantitatively shown to undergo a sharp transition. Based on Fig. 1G of Ref.^33^, *sens* expression drops at 15 ∼ 25 *µ*m away from the DV boundary. Assuming that Wg is secreted by the two stripes of cells with radius 2 *µ*m (based on Fig. 2A from Ref.^44^) at the DV boundary, we set the target gene expression boundary to *x*_b_ = 11 ∼ 21 *µ*m. The overall size of the DV axis patterned by Wg was set to *L* = 70 *µ*m, based on the Hoeschst staining image in Fig. 1E of Ref.^44^.
- **Hh**. The characteristic decay length of Hh was set to λ_Hh_ = 8 *µ*m, as estimated from Fig. S2B of Ref.^41^. The Hh target genes include *en, col*, and *dpp*, whose expression profiles were obtained from Fig. 5 of Ref.^48^. The expression domains of *en, col*, and *dpp*, respectively, span 15 *µ*m, 25 *µ*m, and 30 *µ*m away from the Hh producing cells. Accordingly, the range of boundary positions associated with Hh was set to *x*_b_ = 15 ∼30 *µ*m. The overall size of the domain patterned by Hh was set to *L* = 100 *µ*m, based on Fig. 6C of Ref.^48^.
- **Dpp**. The characteristic decay length of Dpp was set to λ_Dpp_ = 20 *µ*m, as reported in Ref.^5^. The boundary of Dpp target *salm* expression was set to *x*_b_ = 36 ∼ 54 *µ*m, based on Fig. 2H of Ref.^33^. The overall size of the domain patterned by Dpp was set to *L* = 80 *µ*m, based on Fig. 2H of Ref.^33^. Finally, the radius of the sensor for Dpp and Hh mediated patterning was set to *a* = 2 *µ*m, same as for Wg, since they diffuse through the same wing imaginal disk. While we only focus on the boundary defined by *salm* for simplicity, other genes affected by Dpp, including *omb, brk*, and *dad*, also display changes in their spatial expression profiles at locations similar to that of *salm*^33^.

### V. LENGTH SCALES OF CYCLOPS, SQUINT, AND FGF8

In zebrafish embryos, the morphogens Cyclops, Squint, and Fgf8 are responsible for the induction of the endoderm and the mesoderm. All three morphogens are synthesized at the embryonic margin, and diffuse towards the animal pole (Fig. S5a). For simplicity, we will only consider zebrafish embryos around the ∼50% epiboly stage, when the embryo encompasses the top half of the spherical yolk.

- **Cyclops and Squint**. The diffusion coefficient (D) and degradation rate (*k*_d_) of Cyclops and Squint have been measured through ectopic expressions of fluorescently tagged constructs. The resulting decay lengths 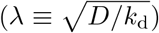 are λ_cyclops_ = 76 *µ*m^75^ and λ_squint_ = 178 *µ*m^75^. The target boundary of Nodal signaling was set by the width of the target genes *gsc* (50 *µ*m) and *ntl* (125 *µ*m) away from the margin as shown in Fig. 1f of Ref.^69^. The width of the domain for Nodal morphogen production was assumed to be 25 *µ*m, as in the kinetic model describing Nodal dynamics in Ref.^69^. The radius of the zebrafish embryo, *R* = 200 *µ*m, was obtained from Fig. 1 of Ref.^74^. Overall, for Nodal morphogens Cyclops and Squint, *z*_b_ = (50*µ*m − 25*µ*m)*/*200*µ*m ∼ (125*µ*m − 25*µ*m)*/*200*µ*m = 0.125 ∼ 0.5.
- **Fgf8**. The characteristic decay length of Fgf8, λ_fgf8_ = 197 *µ*m, was measured from the ectopic expression of fluorescently tagged constructs^78^. Fgf8 affects the expression of target genes such as *spry4, spry2*, and *pea3*. The target boundary of Fgf8 was set by the width of *spry4* (70 *µ*m), *spry2* (85 *µ*m), and *pea3* (120 *µ*m) expression in Fig. 1 of Ref.^74^. Subtracting the width of the Fgf8 expression domain (70 *µ*m) and dividing by the radius of the embryo (200 *µ*m) obtained from the same images lead to *z*_b_ = (70*µ*m − 70*µ*m)*/*200*µ*m ∼ (120*µ*m − 70*µ*m)*/*200*µ*m = 0.0 ∼ 0.25.

### VI .TIME SCALES OF BCD, WG, HH, AND DPP

The expression for the variance of the space-time averaged signal, 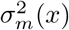 (Eq. 11), is derived with the assumption of a long measurement time, 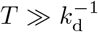 (Appendix 1 of Ref.^19^). Additionally, in order for *Tj*_in_ to represent the total cost of establishing and maintaining the morphogen profile, we require the measurement time to be much longer than the timescale of reaching the steady state (i.e. *T* ≫ *τ* (*x*_b_)). For Bcd, Wg, Hh, and Dpp, *L* ≫ λ and *x*_b_*/*λ ≈ 2, leading to the approximate expression for the characteristic time scale of 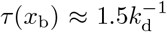 (Eq. 4). We assume that the maximum measurement time, *T*_max_, is set by the time required for the cell to divide, after which the fates of the daughter cells must be re-established.

- **Bcd**.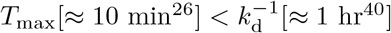.
- **Wg**.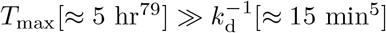.
- **Hh**. 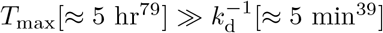.
- **Dpp**.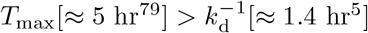.

For Dpp, Hh, and Wg, the space-time averaged measurements may be done for sufficiently long durations required to represent the cost-precision trade-off by *π*_*T*_ (Eq. 14). For Bcd, *π*_*o*_, which is formulated for the point measurement, may better represent the pertinent cost-precision trade-off relation.

**FIG. S4.**
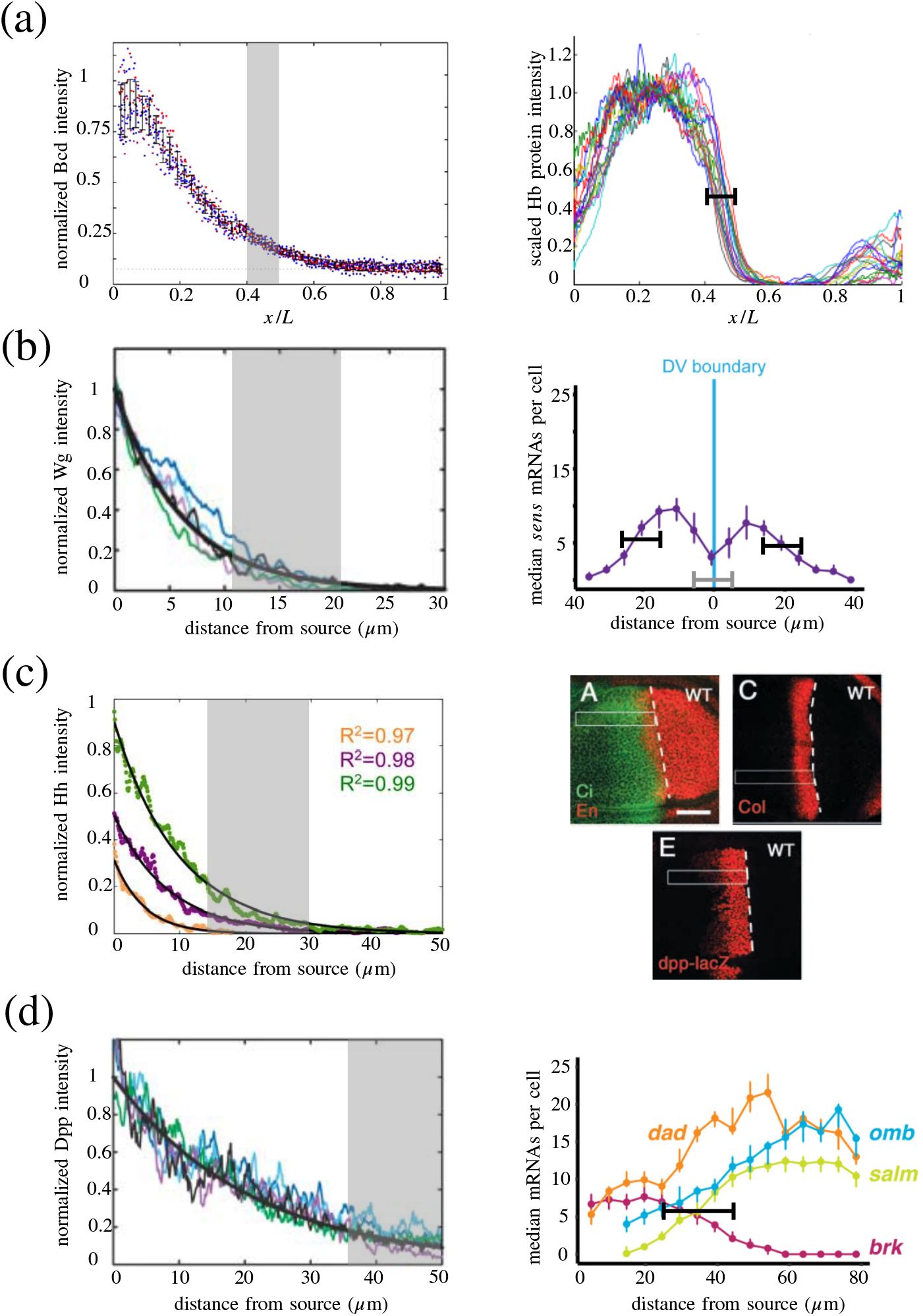
Length scales of naturally occurring morphogen profiles. The left column depicts the exponentially decaying morphogen profiles. The grey shaded regions in the left column depict the respective range of target boundary positions. The right column depicts the target gene expression profiles from which the range of boundary positions are obtained. **(a)** Bcd morphogen profile from Fig. 5a of Ref.^4^. Bcd target Hb protein expression profile from Fig. 3c of Ref.^42^. The black interval denotes the range of boundary positions, *x* = *x*_b_. **(b)** Wg protein expression profile from Fig. 3I of Ref.^5^. Wg target *sens* gene expression profile from Fig. 1g of Ref.^33^. The black intervals denote the range of boundary positions, *x* = *x*_b_. The grey interval denotes the range of cells synthesizing the Wg protein. **(c)** Hh protein expression profile from Fig. S2B of Ref.^41^. The colors green, purple, and orange correspond to different stages of development. The characteristic decay length corresponding to the purple profile, λ ≈ 8 *µ*m, was used for the figures shown in the present work. Hh target Engrailed, Collier, and Dpp protein expression profiles from Fig. 5a, c, and e of Ref.^48^. The white dotted line denotes the AP border, as specified by the staining of Ci. The length of the expression domains of the target genes were estimated from the 50 *µ*m scale bar. **(d)** Dpp protein expression profile from Fig. 1c of Ref.^5^. Dpp target gene expression profiles from Fig. 2h of Ref.^33^. The black interval denotes the range of boundary positions, *x* = *x*_b_.

**FIG. S5.**
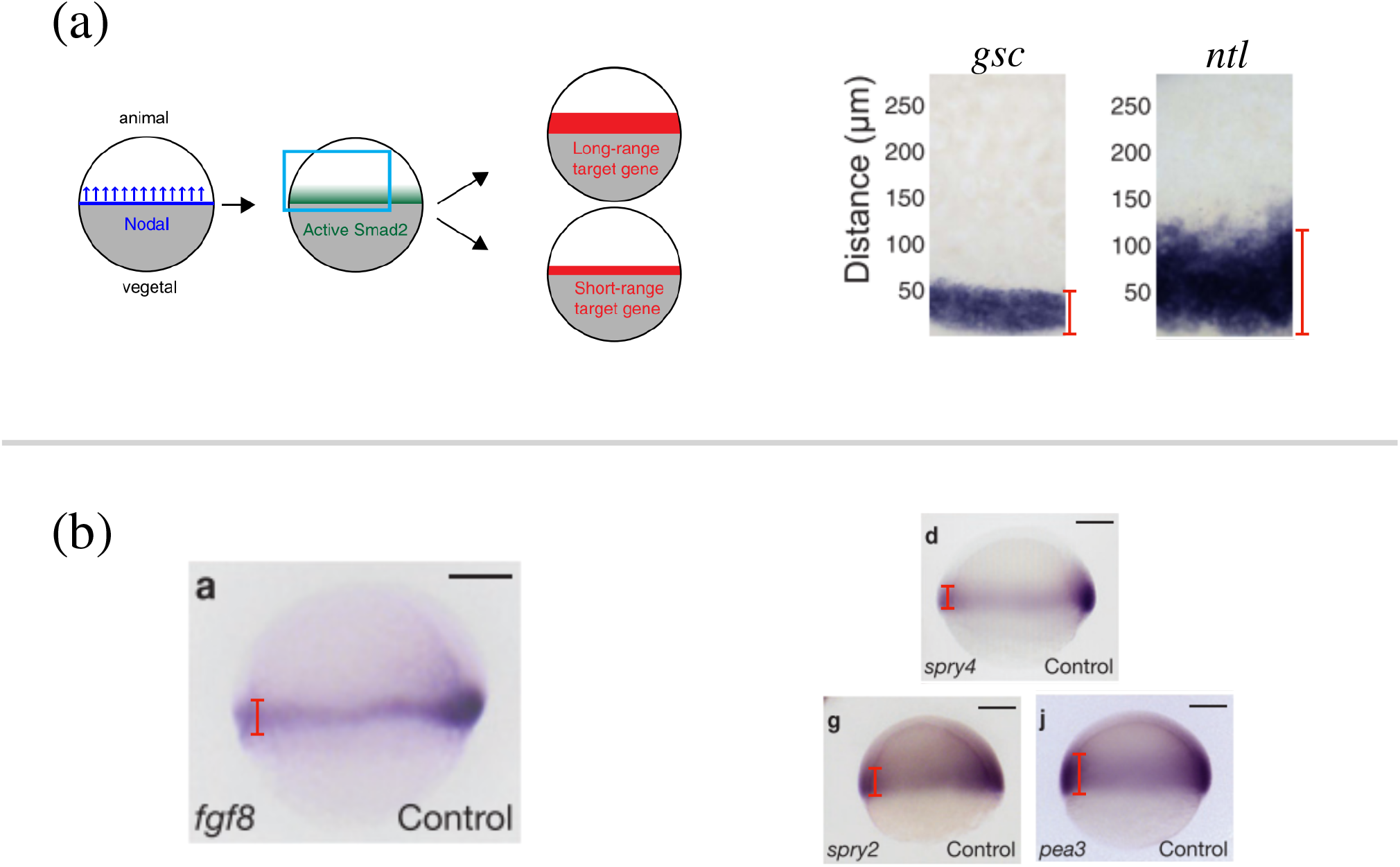
Length scales of naturally occurring morphogen profiles. **a** (Left) Schematic of the Nodal ligands Cyclops and Squint patterning the vegetal-animal axis of the zebrafish embryo. The Nodal signals activate Smad2, which affects the expression of short-range and long-range target genes. (Right) The expression profile of Nodal target genes *gsc* and *ntl* obtained by *in situ* hybridization. The profiles are from zebrafish embryos 5 hours post fertilization, when the embryo covers roughly 50% of the yolk. The red lines denote the width of the target gene expression domain away from the vegetal margin. From Fig. 1 of Ref.^69^. **b** (Left) Expression of *fgf8* in 60% epiboly whole-mount embryos obtained by *in situ* hybridization. Animal side of the embryo is to the top, and dorsal is to the right. (Right) The expression profiles of the Fgf8 target genes *spry4, spry2*, and *pea3*. The red lines denote the width of the target gene expression domain away from the vegetal margin. Scale bars=100 *µ*m. From Fig. 1 of Ref.^74^.

**TABLE S1.** Various length scales associated with the naturally occurring morphogen profiles found in the fruit fly (*) and zebrafish (†) embryos. Further details on how each value was obtained from the associated reference are provided in the SI Appendix *Length scales of Bcd, Wg, Hh, and Dpp* and *Length scales of Cyclops, Squint, and Fgf8*.

| | $L^*$ or $R^\dagger$ ($\mu\text{m}$) | $\lambda[\equiv \sqrt{D/k_d}]$ ($\mu\text{m}$) | $x_b^*$ or $z_b^\dagger$ ($\mu\text{m}$) | $a$ ($\mu\text{m}$) |
| --- | --- | --- | --- | --- |
| Bcd* | 475<sup>4</sup> | 100<sup>4</sup> | 190~238<sup>42</sup> | 2.6<sup>4</sup> |
| Wg* | 70<sup>44</sup> | 6<sup>5</sup> | 11~21<sup>33,44</sup> | 2<sup>44</sup> |
| Hh* | 100<sup>48</sup> | 8<sup>41</sup> | 15~30<sup>48</sup> | 2<sup>44</sup> |
| Dpp* | 80<sup>33</sup> | 20<sup>5</sup> | 36~54<sup>33</sup> | 2<sup>44</sup> |
| Cyclops<sup>†</sup> | 200<sup>74</sup> | 76<sup>69</sup> | 25~100<sup>69</sup> | 10<sup>78</sup> |
| Squint<sup>†</sup> | 200<sup>74</sup> | 178<sup>69</sup> | 25~100<sup>69</sup> | 10<sup>78</sup> |
| Fgf8<sup>†</sup> | 200<sup>74</sup> | 197<sup>78</sup> | 0~50<sup>74</sup> | 10<sup>78</sup> |

